# Fundamental constraints in synchronous muscle limit superfast motor control in vertebrates

**DOI:** 10.1101/135343

**Authors:** AF Mead, N. Osinalde, N. Ørtenblad, J. Nielsen, J. Brewer, M. Vellema, I. Adam, C. Scharff, Y. Song, U. Frandsen, B. Blagoev, I. Kratchmarova, CPH Elemans

## Abstract

Superfast muscles (SFM) are extremely fast synchronous muscles capable of contraction rates up to 250 Hz, enabling precise motor execution at the millisecond time scale. To allow such speed, the archetypal SFM, found in the toadfish swimbladder, has hallmark structural and kinetic adaptations at each step of the conserved excitation-contraction coupling (ECC) pathway. More recently SFM phenotypes have been discovered in most major vertebrate lineages, but it remains unknown whether all SFM share ECC adaptations for speed, and if SFM arose once, or from independent evolutionary events. Here we use genomic analysis to identify the myosin heavy chain genes expressed in bat and songbird SFM to achieve rapid actomyosin crossbridge kinetics and demonstrate that these are evolutionarily and ontologically distinct. Furthermore, by quantifying cellular morphometry and calcium signal transduction combined with force measurements we show that all known SFM share multiple functional adaptations that minimize ECC transduction times. Our results suggest that SFM evolved independently in sound producing organs in ray-finned fish, birds, and mammals, and that SFM phenotypes operate at a maximum operational speed set by fundamental constraints in synchronous muscle. Consequentially, these constraints set a fundamental limit to the maximum speed of fine motor control.

## Introduction

Superfast muscle (SFM) is the fastest known synchronous muscle phenotype in vertebrates^1^. Their ability to repetitively contract and relax fast enough to produce work at cycling rates over 100 Hz and up to 250 Hz sets them apart from other muscles by almost two orders of magnitude and allows the execution of central motor commands with millisecond temporal precision ^1–5^. The phenotype is defined by its mechanical performance and therefore establishing a muscle to be superfast requires quantification of its performance-profile^1^. Although once considered an extremely rare muscle phenotype, SFMs have now been established in most major vertebrate lineages (mammals^5^, birds^3,4^, reptiles^6^, ray-finned fish^6^). SFM in the bat larynx controls the rapid, high frequency calls used by laryngeally echolocating bats to detect, orient to and track prey^5^. SFM in the songbird syrinx controls the precisely timed, rapidly produced acoustic elements and frequency sweeps of bird vocalizations^3,4^ used for territorial defense and mate attraction ^7^. SFM in the Oyster toadfish swimbladder and rattlesnake tailshaker both sets the fundamental frequency of the produced sound by rhythmically contracting the swimbladder and shaking tail rattles respectively ^6^. Other muscles have been suggested to be of the superfast phenotype, such as some extraocular or limb muscles^8^, but non-isometric mechanical tests are lacking to classify them as such. Taken together, SFM seems a commonly occurring muscle phenotype that interestingly so far has been established only in motor systems involving sound production and control.

Force modulation in vertebrate skeletal muscles is precisely timed via the highly conserved excitation-contraction coupling (ECC) pathway that consists of several sequential steps whose individual kinetics affect the maximally attainable force modulation speed. In brief, firing of a motor neuron first triggers the release of intracellular calcium ions (Ca^2+^) stored in the sarcoplasmic reticulum (SR). Subsequent binding of Ca^2+^ to troponin in sarcomeres triggers the exposure of binding sites for the motor protein myosin along actin filaments, allowing the cyclical binding and unbinding of myosin motor head domains to actin, forming actomyosin crossbridges that generate force. Finally, force decreases when SR Ca^2+^-ATPases (SERCA) pump Ca^2+^ back into the SR, consecutively lowering the cytoplasmic free Ca^2+^ concentration ([Ca^2+^]_i_) and returning thin filament inhibition thus preventing further crossbridge formation. Extensive study of SFM in the oyster toadfish (*Opsanus tao*) swimbladder has demonstrated that that no single step in the ECC pathway is rate-limiting, but that multiple hallmark traits have adapted to allow superfast cycling rates. In this fish, a far greater proportion of cellular volume is dedicated to SR, which increases the number and density of SERCA pumps ^9^. Furthermore small, ribbon-like myofibrils ^10^ greatly reduce diffusion distances for Ca^2+^. Together, these adaptations lead to the shortest [Ca^2+^]_i_ transient times observed in any muscle ^6^. Additionally, the rate of actomyosin crossbridge detachment in SFM of the toadfish swimbladder and rattlesnake tailshaker is extremely fast, which is necessary to ensure a rapid force drop after the return of actin filament inhibition ^9^. Whether these hallmark ECC pathway adaptations are shared with avian (songbird syrinx) and mammalian (bat larynx) SFM is currently unknown.

In vertebrates, kinetic tuning of the actomyosin crossbridge cycle for different energetic and biomechanical requirements is accomplished largely via the differential expression of specific myosin heavy chain (*MYH*) genes each with unique kinetic properties ^11^. Because of its thoroughly characterized phylogeny ^12^ and a demonstrated role in force generation ^13^ the MYH gene family is well suited to shed light onto the evolutionary origin and timing of SFM phenotypes. However MYH characterization and expression are currently not known for any SFM. Bat and songbird SFM are excellent models to (i) determine whether SFMs share a common evolutionary origin and to (ii) improve our understanding of muscle function, because sequenced genomes^14^ and identified neural substrates for quantifiable, learned vocalizations exist in some of these taxa ^15–17^.

Interestingly, functional trade-offs associated with the above ECC adaptations do appear to be common to all SFMs. The fast actomyosin detachment rate of SFM in the toadfish swimbladder and rattlesnake tailshaker, in the absence of a commensurate increase in the rate of crossbridge formation, reduces the proportion of myosin motors actively bound to actin at a given time, and thus force and power production ^9^. Furthermore, the increased volumetric allocation for required SR and mitochondria leave less space for contractile machinery, further reducing volume-specific force and power ^18^. Consequentially, the SFMs in swimbladder and tailshaker trade force for speed, constraining movement to very low masses at low efficiency ^9,18,19^. SFM in bat, songbird and toadfish, which produce work up to similar cycling limits (around 200 Hz), develop very similar low force (10-20 kN/m^2^)^1,3–6,20^. Because of the bias for the SFM phenotype to have evolved in sound production systems we asked whether this apparent functional convergence in SFM force profiles is 1) the result of selective pressures common to motor control of sound production and modulation, or 2) due to constraints inherent to the otherwise conserved architecture of synchronous vertebrate sarcomeric muscle, or 3) both?

Here we test if SFM share a common evolutionary origin by characterizing *MYH* gene expression in the SFM of zebra finch (*Taeniopygia guttata*) syrinx and Daubenton’s bat (*Myotis daubentonii*) larynx. The employment of evolutionarily and ontologically distinct *MYH* genes suggests separate evolutionary origins of SFM. Furthermore, by quantifying cellular morphometry and [Ca^2+^]_i_ signal transduction in all known SFM, we show that all SFM share hallmark adaptations in the ECC pathway and have identical intracellular calcium dynamics at their operating temperatures *in vivo*. Our data suggest that SFMs converge at a maximum speed allowed by fundamental constraints in vertebrate synchronous muscle architecture. This implies that motor control of complex acoustic communication is fundamentally limited by synchronous vertebral muscle architecture.

## Results

### Myosin Heavy Chain gene presence in SFM Phenotypes

To test the hypothesis that SFMs in the bat larynx and songbird syrinx share a common evolutionary origin (Fig. 1), we used sequenced genomes of zebra finch^14^ and little brown bat (*Myotis lucifigus*), an echolocating bat in the same genus as Daubenton’s bat, to identify the dominant *MYH* genes expressed in their SFM muscles. Among known *MYH* genes, orthologs of human *MYH13* are prime candidates for superfast motor performance ^21^. In humans and other mammals, *MYH13* and five other *MYH* genes (*MYH8*, *MYH4*, *MYH1*, *MYH2*, *MYH3*) are linked head-to-tail in what is known as the fast/developmental cluster ^12^, which arose from multiple gene duplication events, the first believed to have occurred in an early tetrapod ^22^. Here, we identified the orthologous fast/developmental MYH gene clusters in syntenic regions of zebra finch and little brown bat genomes (Fig. 2a,b). To place putative superfast bird and bat MYH genes in the context of well-characterized vertebrate fast myosin evolution, we additionally used human *MYH13* and the slow/cardiac *MYH7* motor domain sequence to identify predicted *MYH* genes by BLASTp in the large flying fox (*Pteropus vampyrus*) ^23^, chicken (*Gallus gallus*) ^24^, Burmese python (*Python molurus*) ^25^, clawed frog (*Xenopus tropicalis*) ^26^, and torafugu puffer fish (*Takifugu rubripes*) ^27^. Phylogenetic analysis of myosin rod amino acid sequence revealed orthologs of *MYH13* in syntenic genomic regions of all mammalian and avian species included in our analysis corroborating previous studies^12,26^. We also found further lineage-specific expansions of fast/developmental clusters (Fig. 2b), though gene convergence events complicate precise phylogenetic reconstruction of more recently diverging genes ^12^. Some ray-finned fishes, including torafugu, only have a single *MYH* gene in duplicated genomic regions syntenic to the tetrapod fast/developmental cluster ^22^, supporting the view that the duplication event at this locus, which gave rise to an ancestral *MYH13* postdated the tetrapod/ray finned fish last common ancestor. Interestingly, laryngeal muscles of the frog, which power vocalizations in the range of 70Hz, express a *MYH* gene (*MyHC-101d*) that does not appear to have descended from the ancestral fast/developmental locus ^26^. The python possesses an ortholog of *MYH3* (*LOC103063479*) at the expected 5’ end of the cluster, but the presence of a 3’ flanking *MYH13* ortholog is unknown due to the presence of a scaffold end.

**Figure 1.**
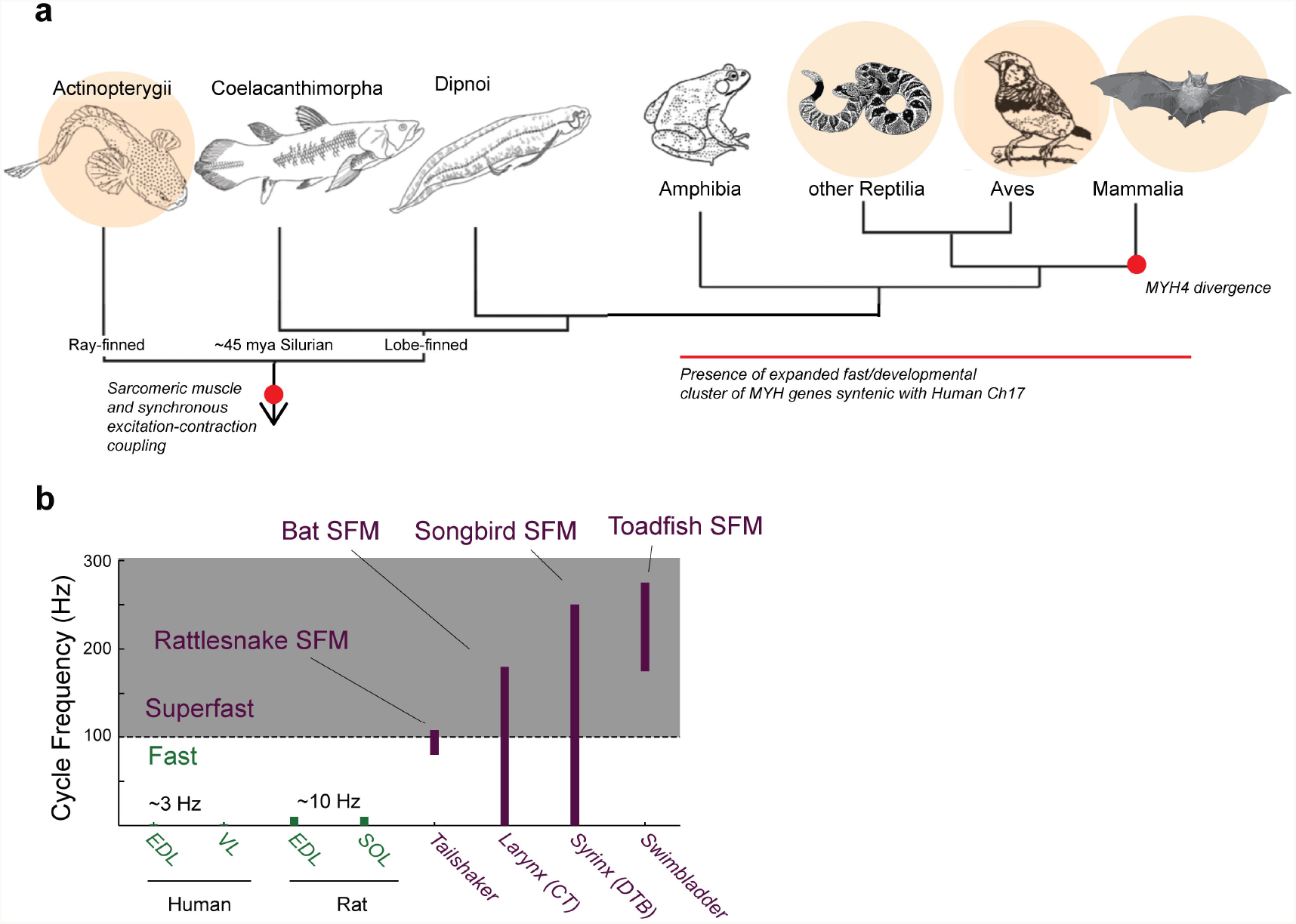
Superfast muscles are present in multiple vertebrate lineages. **a,** Distribution of identified superfast muscles (SFM, beige disks) in vertebrates relative to key evolutionary events: excitation-contraction coupling, and myosin heavy chain (*MYH*) gene evolution. Modified after ^68^. **b,** Ranges of *in vivo* cycling frequency. SFM (labeled purple) has been defined as synchronous muscle capable of producing work at cycling frequencies in excess of 100 Hz (grey box). Bat, songbird, and toadfish SFM approach or exceed 200 Hz. CT, cricothyroid; VTB, *m. tracheobronchialis ventralis*; EDL, *m. extensor digitorum longus*; SOL, *soleus*; VL, *vastus lateralis*.

**Figure 2.**
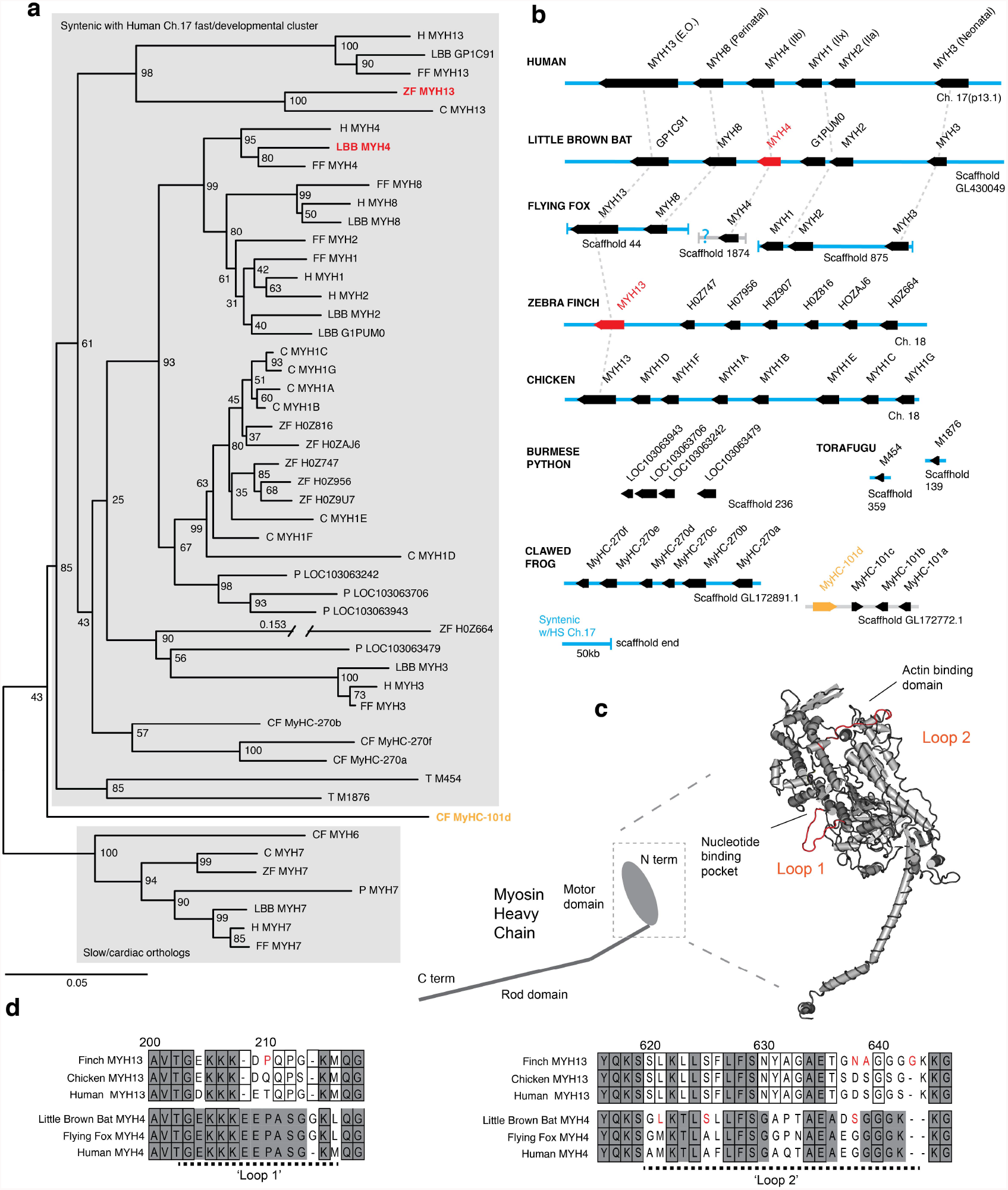
Mammals and avians posess orthologous clusters of myosin heavy chain genes. **a,** Neighbor-joining tree of chicken (C), clawed frog (CF), flying fox (FF), human (H), little brown bat (LBB), Burmese python (P), torafugu (T), and zebra finch (ZF) predicted myosin heavy chain rod domain amino acid sequence from genomic regions syntenic with the human fast/developmental cluster on chromosome 17. Also included are representative cardiac genes, as well as a fast laryngeal myosin gene from the clawed frog (yellow). MYH expressed in bat and finch SFM indicated in red. Human non-muscle *MYH9* was used to root the tree (not shown). Branch lengths shown are derived from a maximum likelihood analysis of aligned amino acid sequence. Bootstrap values from a 1000-replicate analysis are given at nodes in percentages. **b,** Relative local positions of predicted *MYH* genes in genomes of the above species. Synteny with human Ch17 is shown in blue. Red and yellow color-coding as in **a**. Orthologs indicated by vertical dotted lines. **c,** Homology model of human *MYH3* myosin motor domain indicating position of loop subdomains and the nucleotide binding pocket. **d,** Hypervariable surface loops of *MYH13* and *MYH4* motor-domains that likely influence actomyosin crossbridge kinetics. Grey shading indicates conservation with/among *MYH4* orthologs. Outlines indicate conservation with/among *MYH13* orthologs. Substitutions and insertions unique to SFM species in red. Horizontal black dashed lines indicate ‘loop 1’ and ‘loop 2’ subdomains.

To test the hypothesis that *MYH13* orthologs are present in bat laryngeal and zebra finch syringeal SFM, we first used a panel of MYH13-specific commercial and custom-made antibodies (See Methods). Surprisingly, we did not find immunoreactivity against MYH13 in bat laryngeal SFM (Fig. 3a). However a fast myosin antibody (MY-32), which binds gene products of *MYH 1*, *2*, and *4*, showed immunoreactivity. In agreement with that, sequenced amplicons of qPCR analysis revealed that bat SFM laryngeal muscle is highly enriched for the mammal-lineage-specific *MYH4*, which codes for the locomotory MYH also known as MyHC-2B (Fig. 3b). In male zebra finch syrinx SFM, multiple attempts at immunostaining were unsuccessful (see Methods), but sequenced amplicons of qPCR analysis unambiguously showed that it was highly enriched for the *MYH13* ortholog (Fig. 4a). Only one other myosin gene (H0Z747) was expressed and we could not distinguish whether this myosin was expressed in different cells, as suggested by observation of fast myosin antibody (MY-32) staining in starlings ^28^ and zebra finches (**Supplementary Fig. 1**), or co-expressed in the same cells. Extraocular muscle (EOM) expressed *MYH13* at the same level as syringeal SFM (p=0.06, Kruskal Wallis), and also four additional *MYH* genes from the fast/developmental cluster (Fig. 4a).

**Figure 3.**
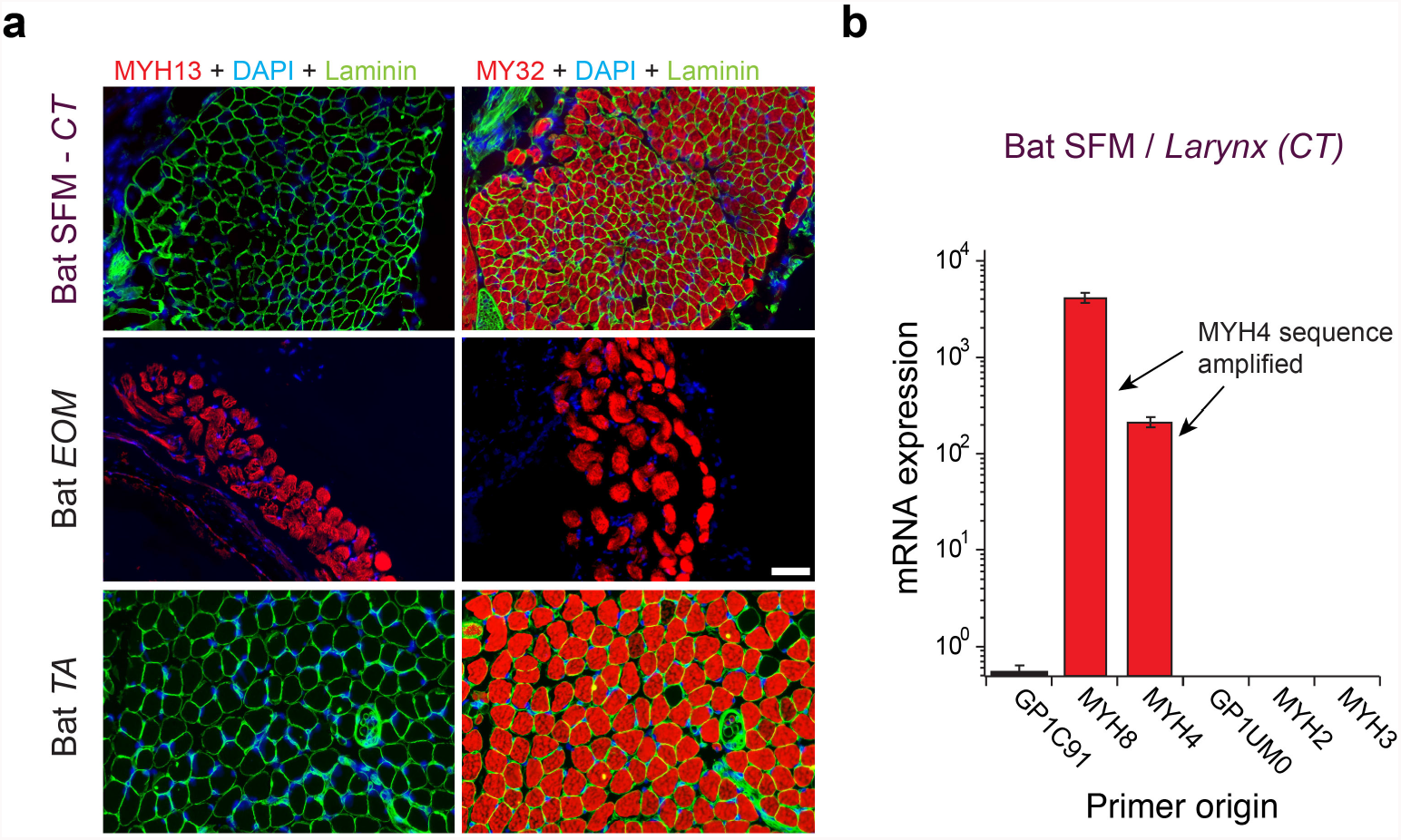
Mammalian superfast muscles are enriched with *MYH4* ortholog. **a** Immunohistochemistry labeling shows absence of MYH13 (left column) in Daubenton’s bat larynx SFM (top) with positive control in extraocular muscle (EOM) (middle) and negative control in body skeletal muscle (*m. tibialis anterior*; TA) (bottom). Fast twitch MYHs (right column) are present in SFM, EOM and TA muscles. **b,** qPCR analysis identifies *MYH4* (MyHC-2b) to be dominating in bat SFM. Both *MYH4* and *MYH8* primer pairs amplified *MYH4* mRNA as confirmed by cDNA sequencing.

**Figure 4.**
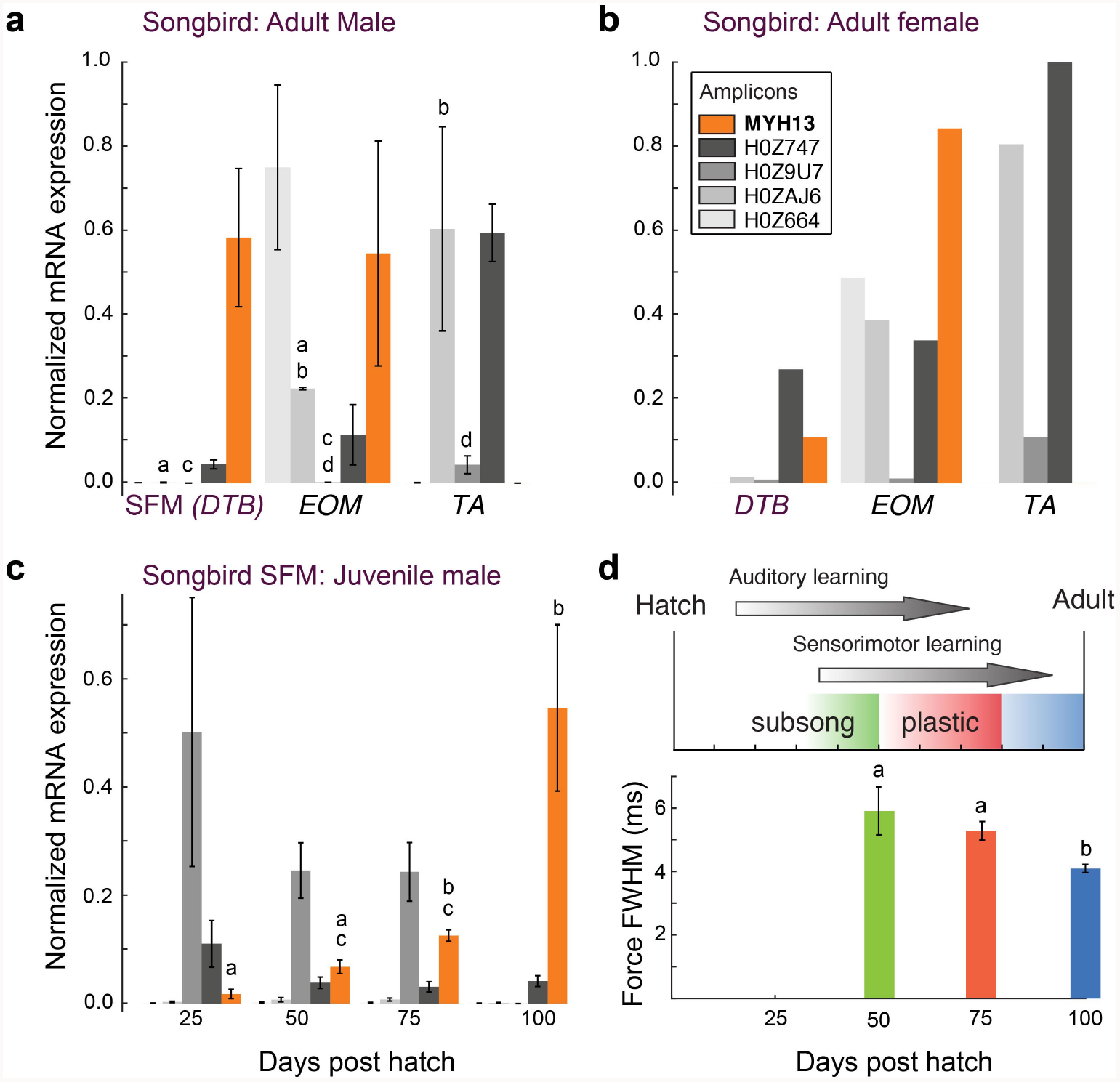
Avian superfast muscles are enriched for the myosin *MYH13* ortholog. **a** qPCR analysis of syringeal SFM of adult male (N=3) and **b,** female (N=3, pooled) zebra finch show high enrichment for the *MYH13* ortholog in singing males. Legend for color code of bars in b. Unshared lowercase letters a-d indicate post-hoc differences with significance level p<0.05. **c,** During the sensorimotor phase of vocal imitation learning in male zebra finches *MYH13* is significantly upregulated (N=3 for each timepoint). Legend for color code of bars in b. Unshared lowercase letters a-c indicate differences with significance level p<0.05. **d,** Syrinx SFM significantly increase their speed during song development (N=4 for each timepoint). FWHM; full width at half maximum.

To identify possible species-specific adaptations to MYH4 in the echolocating bat, and to MYH13 in the zebra finch that could affect crossbridge kinetics, we aligned predicted motor domain amino acid sequence from these genes to their orthologs in the genomes of human, large flying fox and chicken, respectively focusing on two hypervariable regions identified as potential modifiers of crossbridge kinetics (Fig. 2c,d). Loop 1 is situated near the nucleotide binding pocket, and its flexibility is thought to influence the rate of ADP dissociation, and thus is a potentially critical region for the rapid crossbridge detachment seen in SFM^22^. The only sequence difference between little brown bat and human MYH4 in loop 1, an insertion of a glycene at position 214, is also shared with the large flying fox. Although human MYH13 is known to possess rapid ADP dissociation ^21^, the zebra finch ortholog differs from those of both human and chicken by the substitution of a proline at position 210. Loop 2 potentially affects maximum shortening velocity via influence on actin-myosin binding ^29^. Both zebra finch MYH13 and bat MYH4 possess amino acid substitutions in loop 2 and in the case of the finch the insertion of a glycine at position 643 not shared by chicken, flying fox, or human orthologs.

### Associations of MYH Expression and Motor Performance with Singing Behavior

To further explore the relationship between *MYH* gene expression and muscle speed, we next investigated properties of songbird SFM in three groups of zebra finches that differ with respect to song production: juvenile males during the song learning phase singing juvenile, variable song, adult males singing adult, invariable song, and females that do not sing. All three groups produce different types of mostly unlearned calls. Female VTB muscles have two-fold slower twitch kinetics than male VTB muscles ^4^, and expressed more than five times less *MYH13* compared to male VTB muscles (Fig. 4b). Juvenile male songbirds develop their vocal behavior by sensory-guided motor practice in a process bearing many parallels to human speech acquisition ^15,16^. Over the 75 day long sensorimotor phase young zebra finches produce vocalizations that slowly change in acoustical parameters^30^. During this period the variability of acoustic output decreases and accuracy increases, which is attributed to increasingly precise timing of vocal motor pathway activity ^31^. To see whether these vocal changes are associated with changes in SFM performance, we first quantified *MYH* expression in syringeal SFM over song ontogeny. Of all MYH genes, only *MYH13* changed significantly with age (p=0.02, KW-test) and was significantly upregulated progressively (p<0.05, Tukey-Kramer post hoc) in juvenile male syringeal SFM (Fig. 4c). Second, we found that muscle twitch duration significantly decreased progressively (p<0.05, Wilcoxon signed rank tests) during the sensorimotor period of vocal learning (Fig. 4d). Taken together these results suggest that in zebra finch syrinx SFM *MYH13* expression is negatively associated with twitch duration and thus positively with muscle speed.

### Morphometric Cellular Adaptations Converge in SFM

Because the above results established that SFM do not share a common evolutionary origin, we next investigated whether SFM share adaptive mechanisms to ECC pathway traits. We first examined whether cellular adaptations for rapid calcium transients are present across all known SFM. Representative transmission electron-microscopy (TEM) cross-sections of SFM in zebra finch syrinx, bat larynx, toadfish swimbladder and rattlesnake tailshaker showed strikingly higher amounts of SR compared to intraspecific skeletal muscles (Fig. 5a). Based on TEM images we quantified volumetric percentage estimates (VPE) (See Methods) of sarcoplasmatic reticulum (VPE_SR_), mitochondria (VPE_mt_) and myofibrils (VPE_mf_) in all four SFM and intraspecific skeletal muscles (Fig. 5b). Zebra finch syrinx SFM contained 15±1, 22±2 and 50±2% of SR, mitochondria and myofibrils, respectively (n=3) and TEM images resembled those of dove ^32^ and oilbird ^33^ syrinx muscles. Bat larynx SFM contained 24±2, 31±4 and 34±2% of SR, mitochondria and myofibrils, respectively (n=2) (Fig. 5b). SR volume (VPE_SR_) was ≥15% and SR per myofibril ratio (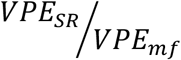 ∗ 100%) was ≥30% for all SFM and significantly higher (p<0.05, KW test) compared to intraspecific skeletal muscles in zebra finch, bat and rattlesnake. Thus the morphometric cellular adaptation of larger SR volume to increase signal transduction^1,18^ is a hallmark adaptation present in all SFM.

**Figure 5.**
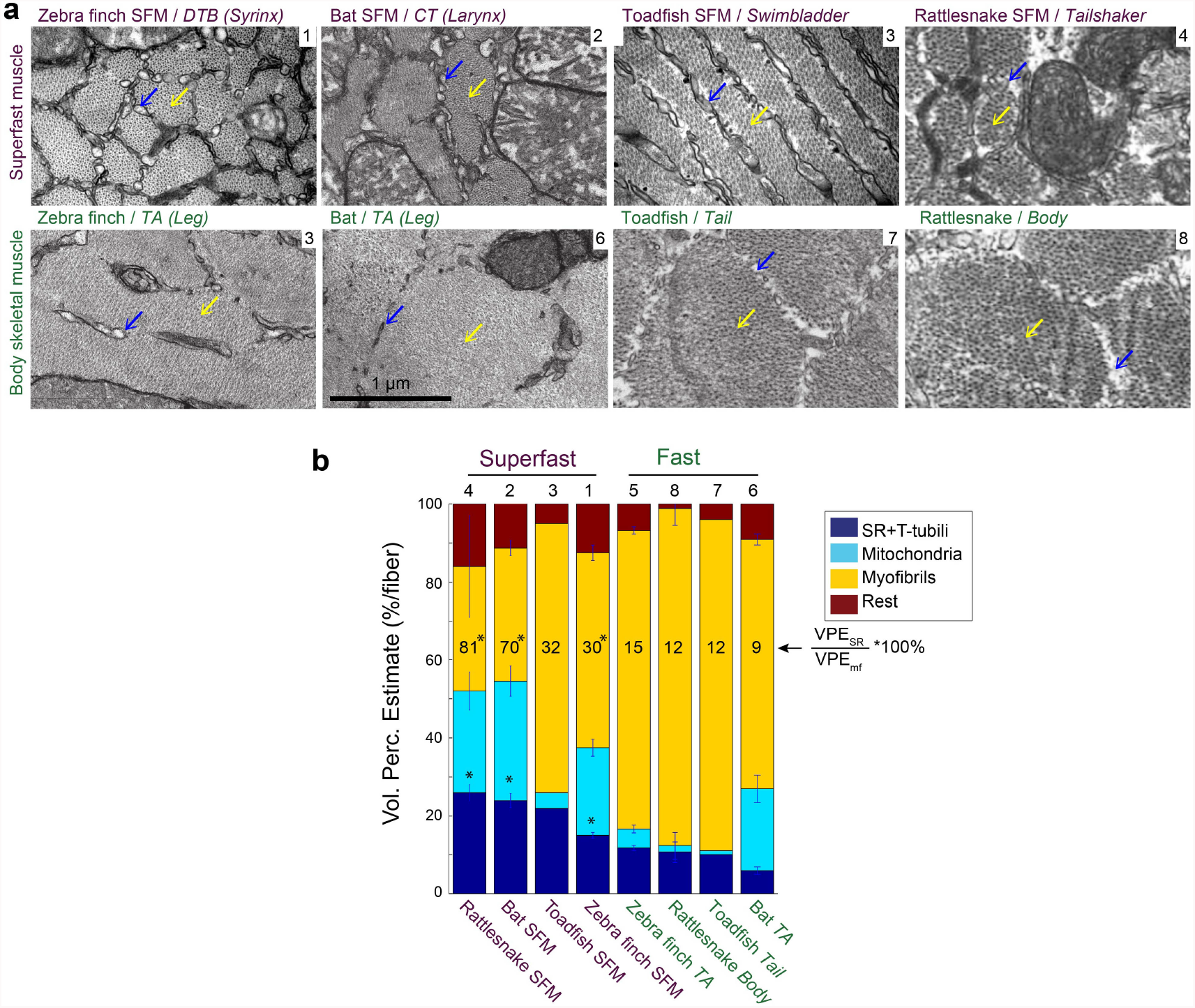
Vertebrate superfast muscles share morphometric cellular adaptations. **a,** Representative transmission electron microscopy images of zebra finch, Daubenton’s bat, Oyster toadfish and Diamondback rattlesnake SFM (upper row) and intraspecific skeletal muscles (lower row). Blue arrows indicate SR, yellow arrows myofibrils. Rattlesnake muscle images adapted from ref 64. **b,** Volumetric percentage estimates (VPE) of myofiber composition (see Methods) show that all SFM have VPE_SR_≥15%, and SR per myofibril ratio ≥30%. Both VPE_SR_ and SR per myofibril ratio are significantly higher compared to fast intraspecific skeletal fibers for zebra finch, bat and rattlesnake (*, p<0.05). Data presented in descending order of SR per myofibril ratio. The numbers 1-8 above the bars refer to images in panel (**a**).

### In vivo [Ca^2+^]_i_ Signal Transduction Dynamics Converge in SFM

We investigated whether [Ca^2+^]_i_ transients in songbird syrinx and bat larynx SFM attain similar extreme kinetics as toadfish swimbladder and rattlesnake tailshaker SFM ^6^. We measured force and real-time [Ca^2+^]_i_ dynamics in zebra finch syrinx and toadfish swimbladder SFM as a function of temperature (Fig. 6). At their respective operating temperatures of 39°C and 25°C, the full width at half maximum (FWHM) values of calcium transients for zebra finch syrinx and toadfish swimbladder SFM did not differ significantly (p=0.54, one-tailed paired t-test) and measured 1.98±0.65 ms (N=4) and 1.91±0.69 ms (N=2), respectively. The FWHM[_Ca2+_] in rattle snake tailshaker SFM is 1.5 ms at 35°C ^6^. Taken together, the FWHM[_Ca2+_] values for SFM in zebra finch, toadfish, and rattlesnake converged to 1.5-2.0 ms at their operating temperatures (Fig. 6b). Because of limited tissue availability of bat laryngeal SFM, we used an *in vitro* assay to quantify calcium uptake and release rates of homogenized SR vesicles as a proxy for whole cell behavior (see Methods). This technique yields comparable values for SFM in zebra finch, toadfish and bat, which were significantly elevated (p<0.01, ANOVA) 5-10 times compared to intraspecific skeletal muscle and also the comparable rat fast twitch *m. extensor digitorum longus* (EDL) (Fig. 6c, **Supplementary Table 1**). These data thus suggest that bat larynx SFM has equally fast calcium dynamics as zebra finch syrinx and toadfish swimbladder SFM. This in turn implies that all established SFM have similar calcium dynamics speeds at their operating temperatures *in vivo*. Concluding, all SFM phenotypes demonstrate hallmark cellular adaptations and functionally convergent calcium transduction dynamics consistent with superfast cycling rates.

**Figure 6.**
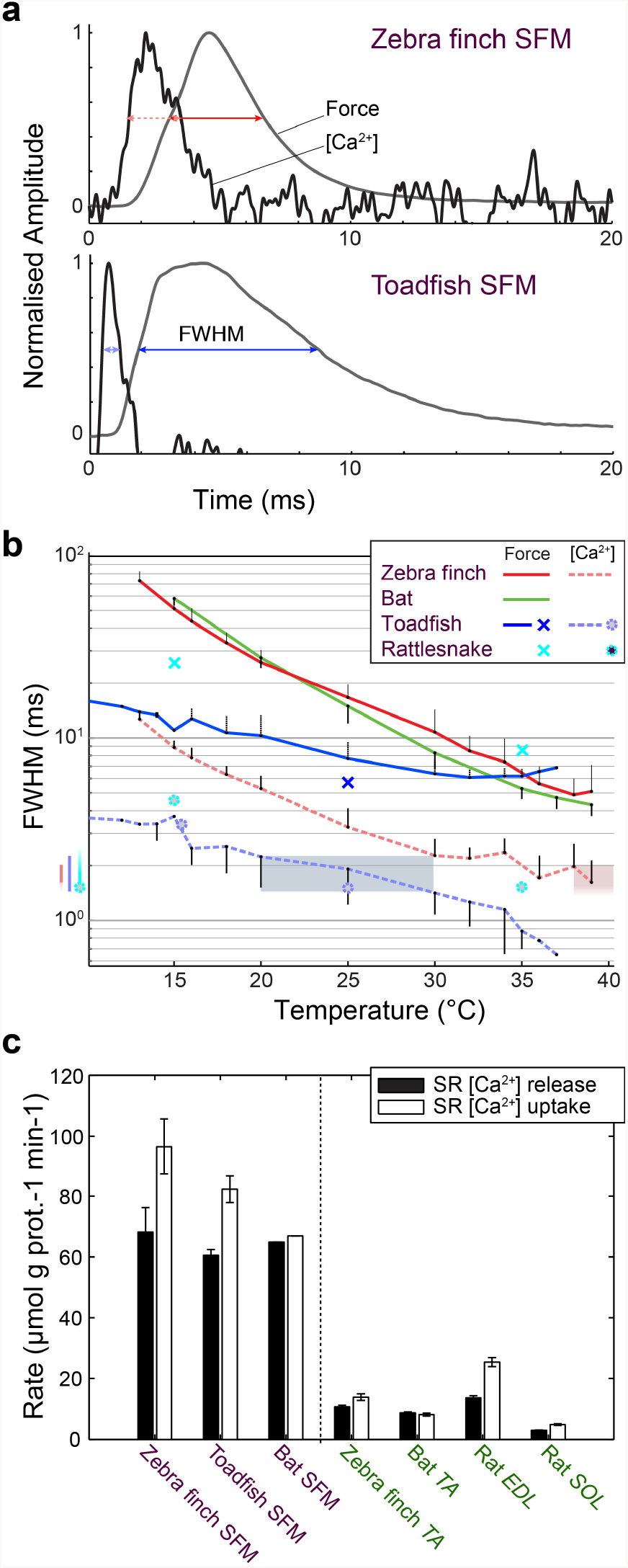
Superfast muscles have convergent elevated myoplasmic calcium transient dynamics. **a,** Representative force development and myoplasmic calcium concentration ([Ca^2+^]_i_) transients in songbird and toadfish SFM. **b,** Force and [Ca^2+^]_i_ transient full-width-at-half-maximum (FWHM) values as a function of temperature. [Ca^2+^]_i_ FWHM values converge to 1.5-2.0 ms for all superfast muscles (color left of ordinate) at their operating temperatures (shaded areas). Filled circle and crosses refer to toadfish and rattlesnake data from ref 7. **c,** Calcium loading and unloading kinetics in isolated SR vesicles (see Methods) yield similar values for bat SFM compared to songbird and toadfish SFM, and were 5-10 times faster than intraspecific skeletal muscle in zebra finch and bat.

## Discussion

We identified the *MYH* genes that power superfast motor performance in songbird syrinx and bat larynx muscle: zebra finch syrinx SFM predominantly expresses the avian ortholog of *MYH13,* which encodes superfast MyHC-sf (aka extraocular MyHC-eo), while Daubenton’s bat larynx SFM expresses an ortholog of the mammalian locomotory *MYH4* (MyHC-2b). Our analysis is consistent with detailed earlier findings ^12,22^ that both *MYH13* and *MYH4* were the result of gene duplication events that occurred in tetrapods, and would, therefore, have no ortholog in the ray-finned fish, the taxon to which the toadfish belongs. Which *MYH* gene is expressed in the rattlesnake tailshaker SFM remains unknown, but no *MYH13* ortholog was found in the Burmese python, which could mean that the gene was lost, or the genome is incomplete and the fast/developmental cluster was truncated by the end of a scaffold. Since extremely fast crossbridge kinetics are necessary, though not sufficient, for superfast performance ^1^, and these are set largely by *MYH* gene expression in vertebrates ^11^, the parsimonious conclusion is that the SFM phenotype found in the sound producing organs of bats, songbirds, and toadfish descend from different evolutionary events. Further support for this notion comes from the observation that these SFM are innervated by motor neurons that run in three different cranial nerves: occipital nerve roots to the swimbladder in fish, the anterior part of the hypoglossus (XII) to the syrinx in birds, and the vagus nerve (X) to the larynx in non-avian tetrapods ^34^.

We further show that all SFM share hallmark ECC pathway adaptations to speed up signal transduction. First, we establish that all SFM converge upon identical speed of *in vivo* calcium dynamics, which can be explained by increased SR per myofibril volume ratio. Additional mechanisms to increase the speed of [Ca^2+^]_i_ transients are also possible and include 1) adaptations that affect the function of ryanodine receptors or the sarcoplasmic reticulum Ca^2+^-ATPase (SERCA) pumps ^35^, 2) faster Ca^2+^ detachment from troponin due to differential expression of TnC genes, or 3) the employment of calcium sequestering proteins such as parvalbumin ^1^. The potential contribution of these mechanisms needs further investigation. Second, actomyosin crossbridge cycling with a higher than normal detachment rate also appears to be a hallmark of SFM, as evidenced by consistently low developed isometric tensions ^1,3–5^. The expression of a *MYH13* ortholog in zebra finch syrinx SFM is consistent with the human protein’s superfast kinetics ^21^, but if and how the observed predicted protein sequence differences between zebra finch and human MYH13 orthologs affect crossbridge kinetics remains unknown. Similarly, further investigation is required to understand how the normally much slower and higher force locomotory MYH4 realizes superfast crossbridge kinetics in bats. The insertion of a second glycine in loop 1 may be a potential mechanism to increase crossbridge cycling by altering mechanical flexibility of the loop. However this insertion is shared by the large flying fox that does not produces calls at high repetition rates, which casts doubt on it being a SFM phenotype specific adaptation. Importantly, superfast crossbridge kinetics may also be the result of adaptations independent of *MYH* genes, so that potential modifiers of crossbridge kinetics, such as myosin light chain genes and phosphorylation state ^36^, also warrant further study.

Both mechanistic as well as functional characteristics are convergent in SFM. Previous observations show that SFM from diverse taxa have remarkably similar maximum performance characteristics (force and cycling frequency) ^1,3,4,6,20,37^. Here we demonstrate that the similarity extends to the speed of signal transduction itself within the ECC pathway (calcium release and reuptake), and to mechanisms of attaining those speeds (volume and distribution of SR). Because we establish SFMs divergent origin, we propose that these similarities must be attributed to convergent evolution. There is good evidence for selective pressures on high cycling speeds in the motor systems in which SFM have evolved: female canaries prefer faster trill rates ^38^, echolocating bats could produce much higher call rates before introducing call-echo ambiguity to their sensory system ^5^ and in toadfish call fundamental frequency, which is set 1:1 by muscle speed ^39^, positively correlates with male fitness ^40^. Thus faster muscles would be expected to evolve.

What mechanisms set the maximum speed to power motion in vertebrates? In toadfish swimbladder SFM a trade-off between force-generating capacity and speed is well-established and results from two constraints related to (i) crossbridge dynamics and (ii) increased ECC pathway requirements ^1^. The consequence of fast relaxation rate is that a lower proportion of myosin motors is bound to actin at a given time reducing the developed force and, by extension, the specific force of the myofibril as a whole ^9^. While detachment rates of intact toadfish SFM fibers ^9^ and isolated MYH13 ^21^ have been shown to be very high, attachment rates are close to that of locomotory muscle, and likely limited by diffusion of the unbound myosin head ^9^. Furthermore, the necessity for very rapid ECC signal transduction requires an increased cell volume dedicated to increased demands on calcium dynamics and ATP production, limiting space for myofibrils, and further diminishing force generation per unit volume of muscle ^18^. Importantly additional SR and mitochondria are likely to affect the muscle cell’s viscoelastic properties. The ultimate limit to how much force can be traded away for speed, while the muscle still maintains the ability to do external work, is dictated by the minimum force required to overcome unavoidable viscous losses associated with shortening. Taken together these observations support the view that SFMs converge at a maximum speed allowed by fundamental constraints in vertebrate synchronous muscle architecture.

Interestingly, only arthropods have pushed the envelope of force/speed tradeoffs further than vertebrate SFM. In insect asynchronous flight muscle force cycling is uncoupled from calcium cycling, minimizing force loss and allowing for high power production at similar cycling speeds to SFM ^41^. However, this increase in power at high speeds is at the great cost of sacrificing precise temporal control of contraction ^41^. The cicada’s synchronous tymbal muscles cycle at over 500 Hz ^42^, but it is unknown if mass reduction or adaptations to the ECC account for this extreme behavior. In the much larger vertebrates, the low forces developed by SFM restrict them to low-mass systems as found in sound production and modulation where precise control is essential.

Myoblast lineage and postnatal motor activation patterns and exercise can play crucial roles in the development of muscle groups or allotypes ^43^ and can alter myosin expression in craniofacial ^44,45^ and body (axial) muscle ^46^. Our discovery of *MYH13* ortholog expression in avian syrinx SFM corroborates earlier developmental studies ^47,48^ and places syringeal myogenic precursor cells in the craniofacial muscle group, because this muscle group can uniquely express *MYH* genes, such as *MYH13*, *15* and *16*, that have never have been found in axial somatic muscle cells. In juvenile songbirds MYH13 expression was progressively upregulated during the sensorimotor period of vocal learning and positively associated with muscle speed (Fig. 4c,d). This observation raises the interesting questions whether *MYH13* upregulation and increase of muscle speed in songbird syrinx SFM is causally related and due to a set developmental program, hormonal influences, muscle exercise in the form of increased neural stimulation, or some combination of these factors. Interestingly the SFM in the larynx of juvenile bats also may increase speed during the first 30 days after birth as suggested by an increase in frequency modulation speed of echolocation calls ^49^. In other motor systems neural stimulation can drive MYH expression patterns: neural activity associated with optokinetic and vestibulo-ocular reflexes stimulates particularly *MYH13* expression in rat EOM ^45,50^, and transnervation from cranial nerve X with XII increased speed of laryngeal muscles in dogs ^51^. Because in EOM *MYH13* is found close to the neuromuscular junction ^52^, a possible mechanism could be that both electrical and chemical activation of the motor neuron directly stimulate *MYH13* production linked to actetylcholine receptors ^53,54^. We speculate that vocal muscle training can be associated with optimizing SFM function. Reversely, our data suggest that, next to neural constraints, SFM can peripherally constrain the execution precision of skilled motor sequences during birdsong and echolocation call ontogeny.

To conclude, our data suggest that SFMs operate at the maximum speed allowed by vertebrate synchronous muscle architecture. In all cases their evolution involved the same specific set of constraints (high cycling speeds and synchronous control) and allowances (minimal force and power), particular to motor systems associated with sound production. Motor-driven acoustic modulation rates in complex communication are thus fundamentally limited by muscle architecture.

## Methods

## Subjects and tissue collection

All procedures were carried out in accordance with the Danish Animal Experiments Inspectorate (Copenhagen, Denmark).

*Songbird:* All bird data presented were collected in zebra finches (*Taeniopygia guttata* (Vieillot, 1817). All adult birds (older than 100 days) were kept in group aviaries at the University of Southern Denmark, Odense, Denmark, and juvenile zebra finches used for muscle kinetics measurements were raised in separate cages together with both parents and siblings at 13 h light:11 h dark photoperiod and given water and food *ad libitum*. The juvenile male zebra finches used for *MYH* mRNA quantification came from the long-term breeding colony at the Freie Universität Berlin, Germany. These animals were kept on a 14 h light:10 h dark photoperiod and given water and food *ad libitum*. We dissected syringeal muscles (*M. tracheobronchialis dorsalis*; DTB, and *tracheobronchialis ventralis*; VTB) on ice immediately after isoflurane euthanasia ^4^. Extraocular (*m. rectus* and *m. oblique*; EOM), flight (*m. pectoralis*; PEC) and leg (*m. tibialis anterior*; TA) muscles were collected as reference tissues.

*Toadfish:* Adult male oyster toadfish (*Opsanus tau,* Linnaeus 1766) were obtained from the Marine Biological Laboratories (MBL, Woods Hole, MA, USA) and housed individually in 80×40×40 cm seawater tanks on constant flow-through of fresh seawater at the Fjord & Bælt field station, Kerteminde, Denmark. Toadfish were euthanized by a blow on the head and double pithing. Superfast swimbladder muscle was isolated bilaterally from the midsection of the swimbladder.

*Bats*: Efforts were focused on Daubenton’s bat (*Myotis daubentoni*) where superfast behavior of the anterior portion of the laryngeal muscle *m. cricothyroideus anterior* (ACTM) was previously established ^5^. We obtained permission (Licenses SNS-3446-00001 & NST-3446-00001) to capture four individuals of *Myotis daubentoni* from the Skovog Naturstyrelsen Inspectorate (Denmark). After isoflurane euthanasia, the CT muscle was isolated and extraocular (EOM) and leg (*m. tibialis anterior*, TA) muscles were collected as reference tissues.

*Rats:* Adult male Sprague Dawley rats (*Rattus norvegicus*) were purchased from the Institute of Biomedicine, Odense University Hospital. The rats were housed in cages with a 12 h:12 h light:dark cycle and provided unrestricted access to water and food. The rats were killed by a blow on the head followed by cervical dislocation. *M. soleus* (SOL) and *m. extensor digitorum longus* (EDL) were collected.

## Myosin heavy chain phylogeny

Fast/developmental myosin heavy chain gene clusters were identified in the genomes of *Myotis lucifigus* and *Taeniopygia guttata* by BLASTP using ENSEMBL genome resources (http://useast.ensembl.org/Myotis_lucifugus/Info/Index and http://useast.ensembl.org/Taeniopygia_guttata/Info/Index) with the rod domain of human *MYH3* (conserved motor domain/rod junction proline 839 to C terminus) as query sequence. Seven Ensembl genebuild annotated predicted proteins were clustered on forward strand of the zebra finch chromosome 18. Six predicted proteins in bat were similarly clustered on the reverse strand of scaffhold GL430049 (**Supplementary Table 2**). The Ensemble annotation for little brown bat *MYH13* was missing exons 1 through 12. Fgenesh gene-finder (hidden markov model based gene structure prediction - http://linux1.softberry.com/berry.phtml?topic=fgenesh&group=programs&subgroup=gfind) was used to partially reconstruct the full-length gene, including loop 1 and loop 2 subdomains, from flanking genomic sequence (Fig. 2). TBLASTN using the same query sequence returned no additional *MYH* domains in these genomic regions.

Fast/developmental cluster and cardiac orthologs from flying fox, chicken, and clawed frog genomes were identified by BLAST using ensemble genome resources, and NCBI genome, with human *MYH3* and *MYH7* (conserved motor domain/rod junction proline 839 to C terminus) as query sequence. Torafugu sequences were obtained directly from ^22^. 2D dot-plot analysis of genomic DNA vs. human MYH3 protein sequence was used to identify clawed frog genes described in^26^, which had been named using scaffold names from an earlier assembly (**Supplementary Table 2**). Fgenesh gene finder was used to reconstruct MyHC-101d sequence from genomic sequence on scaffold 172772.1.

Predicted protein sequences were downloaded and analyzed using Macvector 14.5 software (Macvector Inc. Apex, NC). Human MYH sequences were obtained from the NCBI protein database (**Supplementary Table 2**). To remove the influence of insertions/deletions, multiple protein sequence alignments of human sarcomeric *MYH*s, along with non-muscle *MYH9,* were performed using Clustal W^55^ in Macvector on rod domains from the conserved proline at the motor domain/rod junction (839 in MYH3) to the C terminus.

To reconstruct the phylogeny of fast/developmental cluster *MYH* genes from zebra finch, little brown bat, chicken, large flying fox, Burmese python, clawed frog, torafugu and human genomes^14,23–27^, alignments were performed on 556-residue regions of the rod domain starting with the conserved alanine (1331 in human *MYH3*). Here full-length rods were not aligned due to a region of unknown sequence in bat *MYH13.* Maximum likelihood trees were generated and branch lengths calculated in Macvector using the Neigbor Joining method ^55^ rooted by the non-muscle *MYH9* (Fig. 2a, not shown). Bootstrap analysis from 1000 replicates was used to evaluate internal branches.

For the purposes of comparing relevant hypervariable regions within the motor domain, Clustal W alignments were performed on N-terminal motor domain predicted protein sequence, terminating at the conserved most-c-terminal proline (839 in human *MYH3*) from human *MYH13*, little brown bat and flying fox *MYH4*; chicken and zebra finch *MYH13*. The three-dimensional homology model in Fig. 2c was generated for human MYH3 protein, with scallop myosin head structures (ProteinData Bank code 1kk8) as a template, using the SWISS-MODEL server ^56^ as in ^21^.

## Immunohistochemistry

Muscle tissue was rapidly dissected and frozen in liquid nitrogen after sacrifice. Immunofluorescence staining was carried out on 5 µm thick frozen sections. After initial washing with PBS three times for 5 minutes, sections were incubated for 20 min in a 1% solution of Triton X-100 (Mannheim, Germany) in 0.01 M PBS (Roche, Mannheim, Germany) and rinsed in PBS three times. The sections were then incubated in 10% normal goat serum (Life technologies, MD, USA) for 15 min, and were thereafter incubated with a mouse monoclonal anti-myosin (Skeletal, fast) antibody (clone MY-32, M4276, Sigma-Aldrich, MO, USA), diluted 1:100; a rabbit polyclonal myosin 13/Extraocular myosin (Hinge region) (MP4571, EOM Biosciences, KY, USA), diluted 1:50; and a rat monoclonal anti-Laminin 3 alpha antibody (4H8-2, ab11576, Abcam, MA, USA), diluted 1:500 in PBS with BSA for 60 min at 37°C. The skeletal fast antibody MY-32 binds to MYH1, 2 and 4. To our knowledge no existing antibody can reliably distinguish between these myosins. After incubation with antiserum and PBS wash, a new incubation in 10% normal goat serum followed, after which the sections were incubated in goat anti-mouse IgG1, Human ads-TRITC (1070-03, SouthernBiotech, AL, USA), donkey anti-rabbit IgG-TRITC (711-025-152, Jackson ImmunoResearch, PA, USA) and goat anti-Rat IgG Alexa 488 (ab150157, Abcam, MA, USA) diluted 1:300 for 30 min at 37°C. The sections were thereafter washed in PBS and then mounted in Vectashield Mounting Medium (H-1500) with DAPI (Vector Laboratories, CA, USA). Images were taken using Leica DM6000 at University of Pennsylvania Microscopy Core Facility.

We explored six antibodies to identify MYH13 in zebra finch, but all attempts were unsuccessful. Summarizing, we tried to raise one poly- and one monoclonal antibody against a peptide derived from ortholog-specific loop 2 sequence, tested three commercially available antibodies (of which one was used successfully in bats above), and tested one antibody used successfully in rabbits ^57^. All these antibodies were tested by experienced workers and were either unsuccessful or lacked positive identification (i.e. absence of evidence).

## RNA preparation and qRT-PCR analysis

Muscle tissue was rapidly dissected and frozen in liquid nitrogen after sacrifice. Tissues were homogenized in Trizol using an Ultra-turrax homogenizer and total RNA was extracted according to manufactureŕs instructions (Invitrogen). Reverse transcription was performed by incubating 1 µg of total RNA with 0.25 µg of random hexamers (Amersham Pharmacia Biotech) and 0.9 mM dNTPs at 65 °C for 5 minutes, followed by incubation with 1x First Strand Buffer (Invitrogen), 10 mM DTT and 200 units of Moloney murine leukemia virus reverse transcriptase (Life Technologies) at 37°C for 1 h. For more details see ^58^. Quantitative PCR was performed using 20 µl reactions containing SYBR^®^ Green JumpStart™ Taq ReadyMix™ (Sigma-Aldrich), 5 µl of diluted cDNA, 0.2 µl of reference dye and 150 nM of each primer. Reaction mixtures were preheated at 95 °C for 2 minutes followed by 40 cycles of melting at 95 °C for 15 s, annealing at 60 °C for 45 s, and elongation at 72 °C for 45 s on a Mx3000P qPCR System (Agilent Technologies). The expression levels of targets genes were measured in triplicate and normalized to GAPDH (in birds ^59^) or HPRT (in bats) gene expression. In bat TA and EOM tissues no reference gene (GAPDH, HPRT, tubulin) could be amplified and MYH expression in these tissues could thus not be normalized. The sequences of the primers used are provided in the **Supplementary Tables 3** and **4**. Amplicons were sequenced (Eurofins Genomics, Germany) after all experiments to confirm identity. We only included genes where we had a positive control in the tissues sampled. Data for zebra finches were normalized to the maximum value of all muscles per gene, after which mean+/-SE was calculated. Because of the lower amount of muscle tissue present in females compared to males ^60^, we pooled the tissue of three females prior to tissue homogenization. To compare across juvenile male birds, we collected syringeal tissue of three individuals of the ages 25, 50, 75 and 100 DPH.

## TEM Volumetric percentage estimates (VPE)

The left syringeal muscle VTB and left TA was dissected in three adult male zebra finches. The left laryngeal muscle ACTM and left TA was dissected in two adult Daubenton’s bats. Tissues were fixed with a 2.5 % glutaraldehyde in 0.1 M sodium cacodylate buffer (pH 7.3) for 24h and subsequently rinsed four times in 0.1 M sodium cacodylate buffer. Following rinsing, muscles were post-fixed with 1 % osmium tetroxide (OsO_4_) and 1.5 % potassium ferrocyanide (K_4_Fe(CN)_6_) in 0.1 M sodium cacodylate buffer for 90 min at 4 °C. After post-fixation, muscles were rinsed twice in 0.1 M sodium cacodylate buffer at 4°C, dehydrated through a graded series of alcohol at 4–20°C, infiltrated with graded mixtures of propylene oxide and Epon at 20°C, and embedded in 100 % Epon at 30°C. Ultra-thin (60 nm) sections were cut (Leica Ultracut UCT ultramicrotome) at three depths (separated by 150 µm) and contrasted with uranyl acetate and lead citrate. Sections were examined and photographed at 40,000x magnification in a pre-calibrated transmission electron microscope (Philips EM 208 and a Megaview III FW camera or JEOL-1400 microscope and a Quemesa camera). The VPE_SR_ and VPE_mito_ and VPE_myo_ were determined by point counting using grid sizes of 135, 135 and 300 nm, respectively ^61^. For zebra finch SFM the VPE_SR_ and VPE_myo_ were based on 168 images obtained from 14 fibres in 2 animals, and the VPE_mito_ was based on 288 images obtained from 24 fibres in 3 animals (4,10,10). For zebra finch TA all the estimates were based on 61 images obtained from 5 fibres in 2 animals. For bat CT all the estimates were based on 80 images obtained from 7 fibres in 2 animals, and for bat TA all the estimates were based on 48 images from 4 fibres in 2 animals. The estimated coefficient of error for ratio estimators were 0.05, 0.03 and 0.08 for the estimates of SR, myofibrils and mitochondria, respectively. The rest volume consisted of myoplasm, lipids, nuclei and glycogen particles. TEM images of the Toadfish white muscle body (as prepared in ref. ^62^) for intraspecific comparison to toadfish swimbladder SFM were kindly provided by Dr. C. Franzini-Armstrong. Other values were taken from the literature: *Crotalus atrox* tailshaker and dorsal mid-body muscle ^63^; and *Opsanus tau* swimbladder muscle ^10^ and teleost white muscle ^64^. Meaningful statistical comparison of the toadfish data could not be performed on published mean values in toadfish.

## Force measurements and real-time calcium imaging

Force development curves of zebra finch syringeal VTB muscle were measured at 50 (n=4), 75 (n=4) and 100 DPH (n=4) at 39°C at optimal stimulus amplitude and muscle length as described previously^4,5^. Real-time force development and intracellular calcium concentration kinetics were measured in adult male zebra finches on a custom-built microscope setup. Briefly, the muscle was suspended between a force transducer (Aurora Scientific 400A) and micromanipulator submerged in temperature controlled (accuracy±0.1°C) Ringers solution. This muscle chamber was mounted on an inverted microscope (Nikon eTI, DFA, Glostrup, Denmark). First, force development curves were measured at a range of temperatures (10-39°C) at optimal stimulus amplitude and muscle length. Second, we flushed the sample with 20 µM BTS in Ringers for 20 minutes to inhibit actomyosin interaction and avoid movement artifacts for the subsequent calcium measurements ^65^. Third, to image intracellular calcium, we used a high-affinity calcium dye ^66^ (Mag-Fluo-4, AM stock (ThermoFisher scientific, Slangerup Denmark, # F14201) in DMSO, diluted to 2µM in ringer’s solution) and incubated the samples for 30 minutes at 21°C after which they were rinsed three times with indicator-free medium. A 20x air objective was used to image the sample using a Metal Halide lamp as illumination source and a FITC filter cube (Nikon, DFA, Glostrup, Denmark) to isolate the calcium dye signal. We imaged a small portion of the sample to increase the signal to noise ratio. The resulting signal was passed through a 535±15nm filter (AHF, Germany) detected by a photo multiplier tube (Model R928, Hamamatsu, Denmark) and amplified (model SR560, Stanford Research Systems). The initial temperate series was now repeated at identical stimulation settings. Force, stimulus, and calcium signals were digitized (National Instruments USB6259), aligned on the stimulus onset and analyzed in Matlab. We performed these measurements in zebra finch syringeal VTB muscle (N=5) and toadfish swimbladder muscle (N=2). The bat laryngeal CT muscle’s force data presented in Fig. 6b was collected as part of a previous study ^5^.

## SR vesicle Ca^2+^ uptake and release

Because of limited tissue availability of bat muscle and the rapid experimental rundown of mammalian muscles *in vitro* at relevant physiological temperatures, we could not optimize the real-time calcium imaging technique for bats and used an alternative technique to measure free calcium uptake and release by isolated SR vesicles in muscle homogenate ^67^. Briefly, the SR vesicle oxalate mediated, calcium uptake and release was measured fluorometrically (Ratiomaster RCM, Photon Technology International, Brunswick, NJ) at 37°C using a fluorescent calcium indicator (indo-1). The [Ca^2+^] release and uptake curves were fitted using exponential equations as previously described ^67^.

## Statistics

Data are presented as mean values±1 SE. *MYH13* mRNA levels in juvenile birds were compared using Kruskal-Wallis test, followed by Dunn’s posthoc test. Juvenile twitch times were compared using Kruskal-Wallis test, followed by Wilcoxon signed rank tests. SR volume and SR per myofibril ratio were compared between each SFM and the intraspecific skeletal muscle using a Kruskal-Wallis test. Statistical test results for the rattlesnake TEM data were reported in ref 64. SR vesicle [Ca^2+^] uptake and release data were compared between SFM and intraspecific reference muscle by one-way analysis of variance (ANOVA).

## Acknowledgments

We thank T. Christensen, P. Martensen, and S. Jacobs for technical assistance, the MRC staff at the Marine Biological Laboratories, Woods Hole, and A. Mensinger for supplying toadfish, A. Surlykke, S. Brinkløv, L. Jakobsen for bat access, F.H. Andreade, C. Moncman, N. Rubinstein for discussion and MYH13 antibodies, C. Franzini-Armstrong for toadfish body muscle TEM images, and S. Baylor for calcium imaging advice. J.M. Ratcliffe, and R.J. Schilder commented on the manuscript.

This study was funded by grants from the National Institute of Arthritis and Musculoskeletal and Skin Diseases Training Grant T32 AR-053461 to AFM, Lundbeck Foundation to NO, NØ and BB, Danish National Research Council/Medical Sciences (FSS) to IK, Danish Research Council (FNU) and Carlsberg Foundation to CPHE.

AFM and CPHE conceived of the study. All authors contributed to data acquisition and analysis: myosin phylogeny (AFM), qRT-PCR (NO, IA, BB, KI), calcium handling (NØ, JB, CPHE), transmission electron microscopy (JN), zebra finch ontogeny force measurements (MV, CPHE), tissue preparation (IA, CS), immunohistochemistry (YS). AFM and CPHE wrote the manuscript. All authors read and approved the final manuscript.

The authors declare that they have no competing financial interests. Correspondence and requests for materials should be addressed to CPHE.

